# Reduced emergent character of neural dynamics in patients with a disrupted connectome

**DOI:** 10.1101/2022.06.16.496445

**Authors:** Andrea I. Luppi, Pedro A.M. Mediano, Fernando E. Rosas, Judith Allanson, John D. Pickard, Guy B. Williams, Michael M. Craig, Paola Finoia, Alexander R.D. Peattie, Peter Coppola, David K. Menon, Daniel Bor, Emmanuel A. Stamatakis

**Affiliations:** Division of Anaesthesia, School of Clinical Medicine, University of Cambridge, UK; Department of Clinical Neurosciences, University of Cambridge, Cambridge, UK; Leverhulme Centre for the Future of Intelligence, Cambridge, UK; The Alan Turing Institute, London, UK; Department of Psychology, University of Cambridge, Cambridge, UK; Department of Psychology, Queen Mary University of London, UK; Center for Psychedelic Research, Department of Brain Science, Imperial College London, London, UK; Data Science Institute, Imperial College London, London, UK; Centre for Complexity Science, Imperial College London, London, UK; Center for Eudaimonia and Human Flourishing, University of Oxford, Oxford, UK; Department of Informatics, University of Sussex, Brighton, UK; Department of Neurosciences, Cambridge University Hospitals NHS Foundation, Addenbrooke’s Hospital, Cambridge, UK; Wolfson Brain Imaging Centre, University of Cambridge, Cambridge, UK

## Abstract

High-level brain functions are widely believed to emerge from the orchestrated activity of multiple neural systems. However, lacking a formal definition and practical quantification of emergence for experimental data, neuroscientists have been unable to empirically test this long-standing conjecture. Here we investigate this fundamental question by leveraging a recently proposed framework known as “Integrated Information Decomposition,” which establishes a principled information-theoretic approach to operationalise and quantify emergence in dynamical systems — including the human brain. By analysing functional MRI data, our results show that the emergent and hierarchical character of neural dynamics is significantly diminished in chronically unresponsive patients suffering from severe brain injury. At a functional level, we demonstrate that emergence capacity is positively correlated with the extent of hierarchical organisation in brain activity. Furthermore, by combining computational approaches from network control theory and whole-brain biophysical modelling, we show that the reduced capacity for emergent and hierarchical dynamics in severely brain-injured patients can be mechanistically explained by disruptions in the patients’ structural connectome. Overall, our results suggest that chronic unresponsiveness resulting from severe brain injury may be due to structural impairment of the fundamental neural infrastructures required for brain dynamics to support emergence.

## Introduction

Understanding how brain structure and function give rise to the functioning of the human mind is one of the major open challenges in contemporary neuroscience. In addition to investigating how specific neuroanatomical regions contribute to brain function, it is also useful to study their dependence on highly distributed spatio-temporal patterns of collective activity arising from the complex interactions between multiple neural systems. At its core, this approach builds on the long-standing conjecture that mental activity may be an *emergent phenomenon* arising from the collective activity of neurons in the brain ^1–4^. Unfortunately, so far empirical investigations of this conjecture have been challenging, due at least in part to the absence of means to practically quantify emergence in experimental data. For the same reason, the potential relationship between emergence and functional and structural properties of the human brain still remains to be empirically investigated.

Thanks to recent technical breakthroughs, these questions can now be rigorously brought together under the same conceptual framework, and empirically investigated on neuroimaging data. By leveraging recent developments at the interface between information theory and dynamical systems, the *Integrated Information Decomposition* (ΦID) framework provides the means to conceptualise and quantify emergence in dynamical systems ^5^. Specifically, it can be rigorously shown that a system exhibits *causal emergence* to the extent that its state as a whole provides information about its future state that cannot be obtained from the states of its individual components ^1, 4^. In other words, causal emergence is *the causal (predictive) power of the macroscale, above and beyond the microscale effects* ^1^. ΦID is widely applicable, offering rigorous methods to reason about and quantify causal emergence across a variety of different systems — from flocks of birds and Conway’s celebrated Game of Life, to functional MRI measurements of human and non-human primate brain dynamics ^1, 6^.

By making emergence in the brain formally quantifiable, ΦID also makes it possible to contextualise how emergence relates to other fundamental properties of brain functional and structural organisation. Conceptually, emergence is deeply intertwined with the idea of *hierarchical organisation,* which (in its many possible conceptualisations ^7^) is a fundamental principle in our contemporary understanding of the brain ^8–17^. Thanks to ΦID, we are now in a position to characterise the empirical relationship between emergence and hierarchy in the activity the human brain.

Physiologically, the conditions for the dynamics of functional brain activity to exhibit hierarchical and potentially emergent properties are shaped by the structural connectome on which they unfold ^18–22^. Therefore, different physical configurations of the structural network may be expected to support different degrees of emergent or hierarchical dynamics. One attractive avenue to tackle the relationship between structural organisation and emergent dynamics is via the recent framework of *network control theory* ^23^, which studies how the organisation of a structural network shapes its ability to influence the functional dynamics that take place over it.

A unique opportunity to investigate how emergence is related to both functional and structural characteristics of the human brain comes from studying patients with chronic disorders of consciousness (DOCs) as a result of severe brain injury. Chronic DOCs involve permanent neuroanatomical damage, including disruption of the brain’s structural connectivity and dynamics ^24–35^. In addition to providing a powerful avenue to relate brain organisation and (dys)function, this approach also addresses a pressing need to understand how the structural and functional brain reorganisation induced by DOC patients’ injuries prevent them from recovering ^36, 37^. Therefore, in the present work we combine functional and diffusion MRI data to study brain function and structure in a cohort of 21 DOC patients and 18 healthy controls. We leverage ΦID and network control theory to investigate the relationship between emergence in brain dynamics, on one hand, and healthy and pathological aspects of the brain’s structural and functional architecture, on the other.

Our main hypothesis was that emergent and hierarchical character of brain activity should be diminished in the brains of severely brain-injured unresponsive patients. Further, we hypothesised that the capability of these patients’ anatomical connectomes to control brain activity should be compromised as a result of their injury. Crucially, these hypotheses are tightly interconnected: emergence and hierarchy are two distinct but complementary ways of viewing the same dynamics, and a controllability shapes the repertoire of dynamics that the structural connectome can entertain. Therefore, as our final hypothesis we expect that causal emergence, functional hierarchy and structural controllability should be related to each other. To obtain mechanistic insights beyond pure correlation, we address this last hypothesis using *whole-brain computational models*, which simulate neurobiologically realistic brain dynamics based on different empirical connectomes ^15, 37–43^. The model-generated dynamics can then be directly interrogated in terms of causal emergence via ΦID, through the same process as the empirical brain dynamics. This approach enables us to seek a mechanistic interpretation of our results. Through these convergent, multimodal investigations we shed light on how healthy and pathological brain structure influences brain dynamics.

## Results

Here, we adopted the recently developed mathematical framework of Integrated Information Decomposition (ΦID) to quantify causal emergence in the dynamics of the human blood-oxygen-level dependent (BOLD) signal from fMRI data of N=18 healthy controls and N=21 DOC patients, further subdivided into N=10 patients diagnosed with unresponsiveness wakefulness syndrome (UWS, also known as the vegetative state), and N=11 patients in a minimally conscious state (MCS), who can occasionally exhibit behavioural signs consistent with transitory responsiveness. Through this powerful new approach to quantify emergence, we sought to investigate the fundamental connection between emergence and human consciousness, and how they both relate to relevant aspects of brain function (spatiotemporal hierarchy) and structure (network controllability).

### Diminished emergence in the brain dynamics of DOC patients

To empirically investigate the hypothesis that the macroscale capacity for emergence is diminished in chronically unresponsive brain-injured patients, we adopted the account of causal emergence recently formalised by ΦID (Methods). A macroscale feature *V_t_* is said to be causally emergent if it has “unique” predictive power over the future evolution of the system *X_t_* — in the sense of providing information about the dynamics of the system that cannot be found in any of the parts of the system when considered separately. Thus, supervenience is a relationship between the macroscale (for example, the shape of a flock of birds) and the microscale (the individual birds) at a particular point in time, whereas emergence pertains to the joint dynamics of the macro- and the microscale ^1^. Crucially, ΦID allows one to measure the maximal amount of unique predictive power that any emergent feature could have with respect to the microscale, which upper-bounds the ability of the system to host emergent features — hence termed “emergence capacity” (see Methods).

Here, we employed ΦID to measure the capability for causal emergence of the coevolving activity of pairs of brain regions, based on their fMRI BOLD signals at rest (see Methods for details of how ΦID’s information-theoretic quantities are computed). In other words, we quantify the capacity of pairs of regions to give rise to emergent behaviour together. By averaging the resulting estimates of emergence capacity across all pairs, we obtained an estimate of the global emergence capacity across the brain for each subject.

An analysis of variance revealed a significant effect of disorder severity (control, MCS or UWS) on the mean values of emergence capacity (F(2,37) = 26.08, *p <* 0.001), with subsequent post-hoc tests (corrected for multiple comparisons using the Benjamini-Hochberg procedure to control the false discovery rate ^44^) indicating that healthy controls had significantly higher capacity for causal emergence than both MCS and UWS patients across brain regions - as well as a trend towards significance for the difference between patient groups (*p =* 0.072) (see Figure 1 and Table S1). Thus, supporting our first hypothesis, we identified that lower causal emergence is observed in chronically unresponsive patients after severe brain injury.

**Figure 1.**
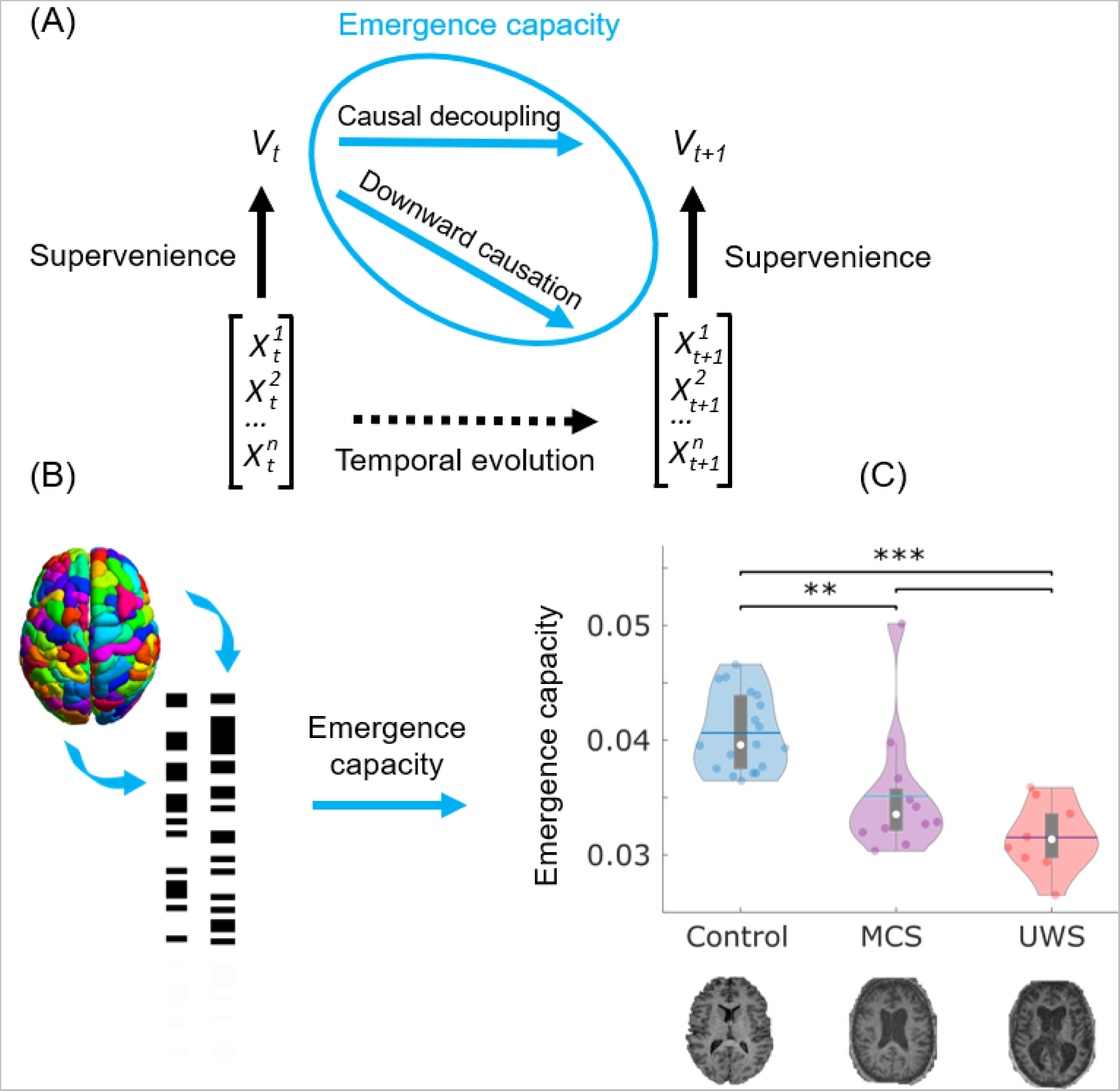
Causal emergence is diminished in the brain dynamics of DOC patients. (A) C) Relationship between emergence and supervenience, illustrating how causal decoupling and downward causation are two different types of emergent phenomena. (B) The global emergence capacity of the human brain is obtained from Integrated Information Decomposition as the average emergence capacity between each pair of discretised regional fMRI BOLD signals (Methods and Figure S1). The framework rests on the following definitions. Given a system composed of multiple elements that co-evolve over time, we say that a macroscale feature 𝑉_𝑡_ is *supervenient* on the state of the system at time *t,* denoted by 𝑋_𝑡_, if 𝑉_𝑡_ is fully determined by 𝑋_𝑡_ (beyond the addition of noise), if anything about 𝑉_𝑡_ that can be predicted from the system’s previous state, 𝑋_𝑡−1_ can also be predicted from the system’s current state, 𝑋_𝑡_(A). Then, a supervenient feature 𝑉_𝑡_ is said to be causally emergent if it has “unique” predictive power over the future evolution of 𝑋_𝑡_ — in the sense of providing information about the dynamics of the system that cannot be found in any of the parts of the system when considered separately. (C) Violin plots of each subject’s emergence capacity by group. Data points represent subjects. White circle, median; centre line, mean; box limits, upper and lower quartiles; whiskers, 1.5x interquartile range. ** p < 0.01; *** p < 0.001, FDR-corrected. We also show that analogous results are obtained using continuous (rather than discretised) signals (Figure S2A), and using a different information-theoretic formalism (Methods and Figure S2B), with UWS patients exhibiting significantly lower emergence capacity than healthy controls in both cases.

### Compromised spatiotemporal brain hierarchy in DOC patients

Emergence is conceptually intertwined with another central concept in the modern neuroscientific literature: hierarchical organisation. In particular, recent theoretical and empirical work has shown that the global activation patterns that arise in response to spontaneous local activity induce a spatiotemporal hierarchy, whereby different regions vary in their capability to elicit spatially distributed neural activity over time — a phenomenon dubbed “intrinsic-driven ignition” ^14^. Crucially, in previous work, this spatio-temporal hierarchy (meaning, roughly, the difference in elicited activity between the most and least influencing regions throughout the brain) was diminished during the transient unresponsiveness induced by both sleep and anaesthesia ^45, 46^. Therefore, having identified an association between diminished emergence capacity and disorders of consciousness due to severe brain injury, we proceeded to test whether DOCs also induce a reduction in the spatio-temporal hierarchy of brain function.

Operationally, intrinsic-driven ignition (IDI) is obtained by identifying “driver events” of unusually high activity in spontaneous BOLD signals of each region and measuring the concomitant activity occurring in the rest of the brain. Importantly, regions generally vary in the extent of the ignition they typically elicit, and the spatial variability of the mean IDI across regions defines the brain’s *spatio-temporal hierarchy*. In other words, when driver events in some regions are able to recruit a large fraction of the brain while events in others not at all, brain dynamics can be characterised as being highly hierarchical ^14, 45^.

Results supported our hypothesis of diminished spatio-temporal hierarchy in the functional brain activity of DOC patients: an ANOVA revealed a significant effect of diagnosis on the spatiotemporal hierarchy of ignition (F(2,37) = 11.28, *p <* 0.001), with follow-up t-tests indicating that UWS patients exhibited reduced hierarchical organisation compared with both MCS patients and healthy controls (Figure 2B and Table S2). Therefore, our results are in line with previous studies on sleep and anaesthesia ^45, 46^, indicating that spatiotemporal hierarchy of the brain’s intrinsic-driven ignition is compromised following the kind of severe brain injury that results in chronic disorders of consciousness.

**Figure 2.**
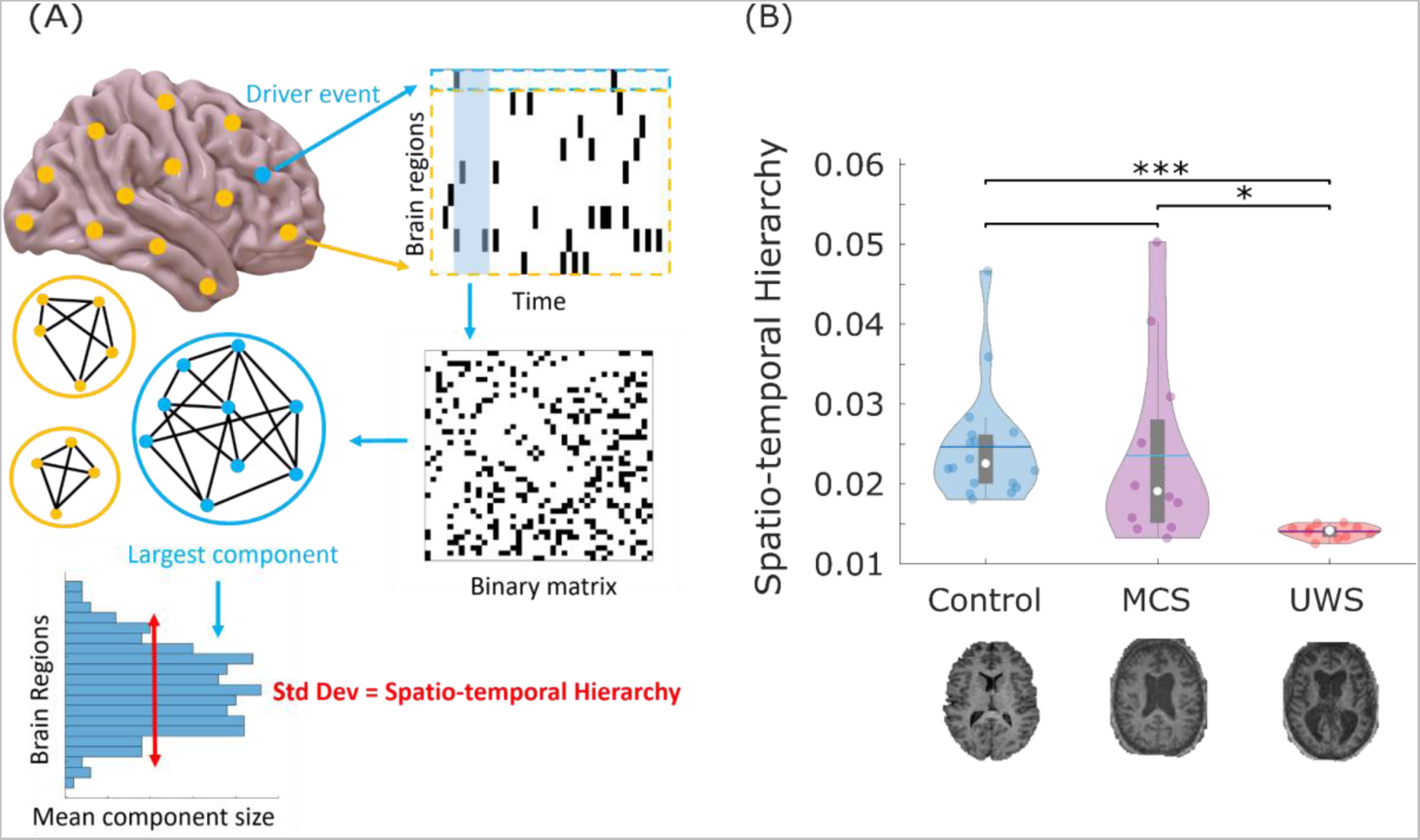
Spatio-temporal hierarchy of intrinsic-driven ignition is compromised in DOC patients. (A) intrinsic-driven ignition is obtained by identifying “driver events” (unusually high BOLD spontaneous activity; here, an event is defined to occur at a given region when its BOLD signal exhibits a Z-score larger than 1, following previous work ^14, 45^), and measuring the magnitude of the concomitant activity occurring in the rest of the brain within a short time window (here, 4 TRs, approximately corresponding to the duration of the hemodynamic response function, following previous work ^14, 45^). The level of intrinsic-driven ignition is calculated as the size of the resulting largest connected component over a network linking regions that exhibit co-occurring events within the chosen time window. A measure of spatio-temporal hierarchy is obtained by calculating the variability across regions of their average IDI. (B) Violin plots of each subject’s spatio-temporal hierarchy by group, showing that UWS patients exhibit diminished hierarchy compared with both healthy controls and MCS patients. Data points represent subjects. White circle, median; center line, mean; box limits, upper and lower quartiles; whiskers, 1.5x interquartile range. * p < 0.05; ** p < 0.01, FDR-corrected.

To ensure the robustness of our results, we repeated our analyses pertaining to both emergence capacity and spatio-temporal hierarchy after controlling for mean framewise displacement as a covariate of no interest (Figure S3) and using a different parcellation size (129 ROIs; Figure S4).

### Reduced network controllability of the DOC connectome

Our results so far have shown that the brain activity of chronically unresponsive brain-injured patients compared to healthy controls is characterised by decreased causal emergence, and, possibly closely related to this, a diminished spatio-temporal hierarchy of brain dynamics. Crucially, however, brain dynamics are fundamentally shaped by the underlying structural connectome on which they unfold ^18–22^ - and indeed DOC patients often exhibit disrupted structural connectivity due to their injury, as well as subsequent complications and atrophy. To study how reductions in emergence and spatio-temporal hierarchy are related to brain structure we leverage principles of *network control theory*, which has recently become a prominent approach to investigate the relationship between the brain’s network structure and its ability to support different kinds of functional dynamics ^23, 47–55^.

Given a system of active elements (e.g., brain regions) interconnected by a network of structural connections (here, from the human connectome project), the organisation of the network’s connections can be studied via control theory to determine how to intervene on the system to achieve a desired configuration of activity of its elements (Figure 3A,B). Specifically, if energy is injected into the system via a particular node or set of nodes, it will spread to the rest of the system according to the network’s connectivity, so that the activity of individual elements will be differently affected. As a consequence, a specific desired pattern of activity may be best achieved by intervening on some nodes rather than others. Nodes requiring comparatively less effort (i.e. smaller input energy) to achieve the same target configuration are said to be more *controllable*. Importantly, network control theory makes it possible to theoretically estimate controllability based on the structural network itself, without the need for physical interventions.

**Figure 3.**
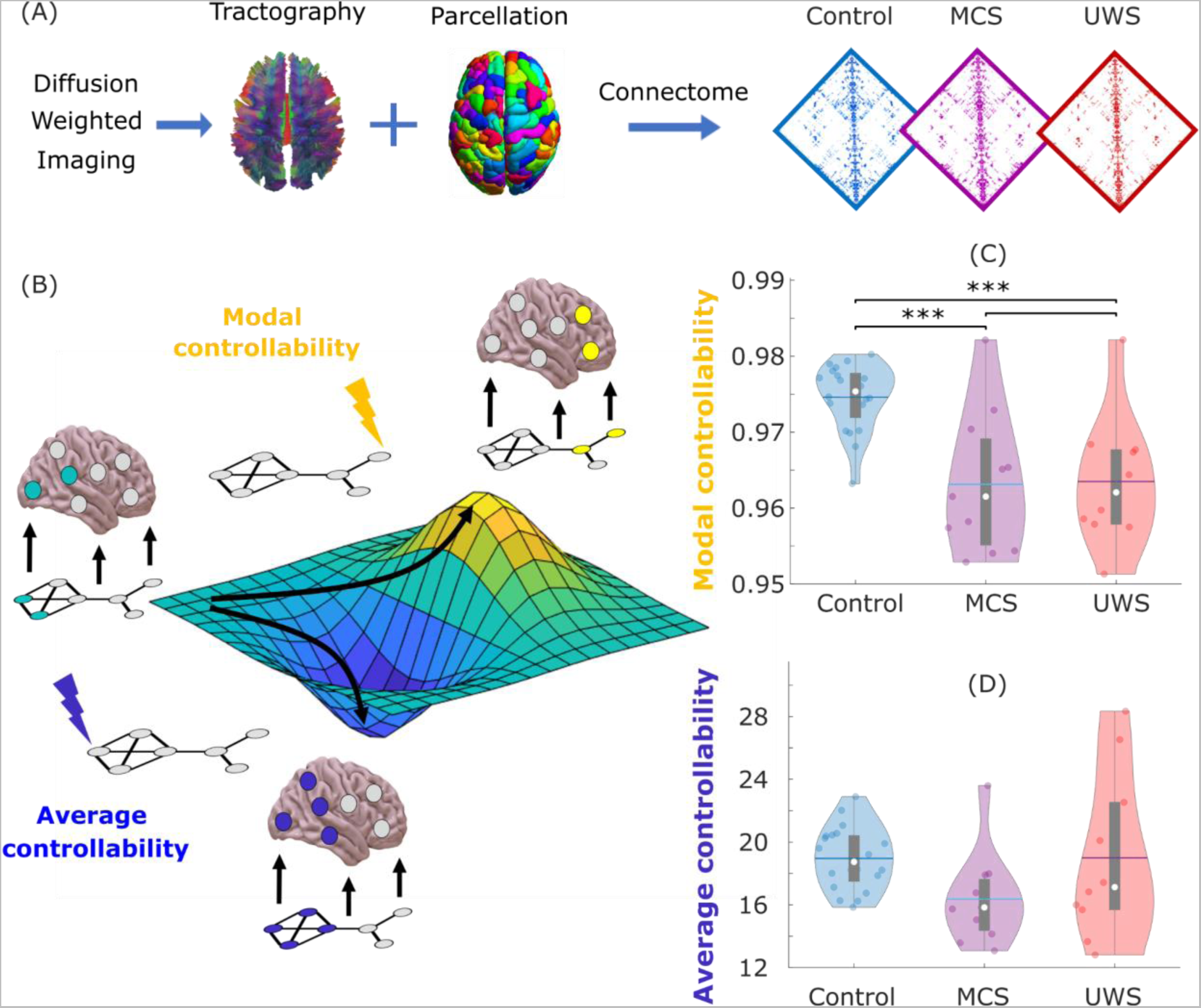
Reduced controllability of structural brain networks in DOC patients. (A) To obtain the structural connectome, diffusion weighted imaging (which measures the direction of water diffusion in the brain) is used to reconstruct white matter streamlines through tractography algorithms, obtaining a network representation of the physical connections between brain regions (here, *N =* 234 regions from the Lausanne atlas). The average structural networks for each group (control, MCS and UWS) are shown. (B) Functional brain activity (colored nodes are active, grey nodes are inactive) evolves through time over a fixed network structure (displayed below the brains). From a given starting configuration of activity (green), some alternative configurations are relatively easy to reach in the space of possible configurations (valley, in blue), whereas others are relatively difficult to achieve (peak, in yellow). To achieve a desired target configuration, input energy (represented by the lightning bolt icons) can be injected locally into the system, and it will spread to the rest of the system based on its network organisation. Average controllability quantifies the network’s support for moving the system from an initial configuration of activity (green) to easy-to-reach configurations (blue), whereas modal controllability quantifies the network’s support for moving the system to difficult-to-reach configurations of activity (yellow). (C) Global average controllability across each group. (D) Global modal controllability is significantly reduced in DOC patients. Data points represent subjects. White circle, median; center line, mean; box limits, upper and lower quartiles; whiskers, 1.5x interquartile range. *** p < 0.001, FDR-corrected.

According to this formalism, different types of controllability can be defined depending on the type of desired outcomes. Here, we focus on two widely adopted and complementary notions: *average* and *modal controllability* (Methods) ^23^. Average controllability refers to the ability to steer the dynamics of the system towards configurations of activity that are relatively easy to reach, in the sense that they would require little energy to be injected into the system, because they are relatively close to the initial pattern of activity (Figure 3B, blue). In contrast, modal controllability refers to steering the dynamics of the system towards patterns of activity that are relatively difficult to reach, because they are very different from the current activity of the system (Figure 3B, red). Considering the widespread alterations typically observed in DOC patients’ brain dynamics ^26, 27, 32, 33, 56–58^, our third hypothesis was that DOC patients should exhibit compromised controllability of their structural connectomes, reflecting a diminished capacity to control brain dynamics towards desired functional configurations.

We used diffusion MRI data to construct a network of structural connectivity for each subject in terms of the number of white matter streamlines connecting each pair of 234 cortical and subcortical regions ^59^ (Methods) (Figure 3A). Based on each individual’s structural connectome, we derived the average and modal controllability of each subject by taking the mean controllability across brain regions ^52^. Analysis of variance revealed a significant effect of diagnosis on whole-brain modal controllability between the three groups (F(2,36) = 14.50, *p <* 0.001) (Figure 3C), while showing no significant differences in average controllability (F(2,36) = 2.79, *p =* 0.075) (Figure 3D). Post-hoc pairwise t-tests (FDR-controlled) to explore the significant effect from the ANOVA indicated significantly higher modal controllability across brain regions for healthy controls than either MCS or UWS patients (Table S4). Analogous results were also obtained when using a different parcellation size (129 ROIs; Figure S5). Thus, our results indicate that the structural connectomes of DOC patients are significantly less suitable to steer their dynamics specifically towards hard-to-reach configurations - in line with existing results about the central importance of appropriate dynamics to support consciousness in humans and other mammals ^26, 27, 60–63^

### Convergent evidence for structural-functional relationships

So far we have demonstrated that the brains of chronically unresponsive brain-injured patients are characterised by reduced emergence and spatiotemporal hierarchy of brain activity, as well as structural network differences, specifically in terms of compromised modal controllability. This set of results raises the question of how exactly functional and structural deficits observed in these brain-injured patients are related with each other.

This section and the next will investigate the structure-function relationship via two convergent approaches: First, by correlating the functional (emergence, spatiotemporal hierarchy) and structural (modal controllability) measures that had exhibited significant differences between controls and DOC patients across all subjects (patients and controls); Second, to derive mechanistic insights beyond correlation, by using whole-brain computational modelling to generate biophysically realistic macroscale dynamics based on different connectomes, thereby illuminating how connectome structure shapes emergence and hierarchy. We report the results of these two investigations in turn below.

Results of the correlation analysis supported our hypothesis, indicating significant positive values of Spearman correlation between all measures: causal emergence, spatiotemporal hierarchy, and overall modal controllability of the structural connectome (Figure 4). These correlations have two key consequences: they support the theoretical link between emergence and spatiotemporal hierarchy, and they confirm our expectation that the presence of emergent and hierarchical dynamics in functional brain activity is related to controllability of the underlying structural connectome.

**Figure 4.**
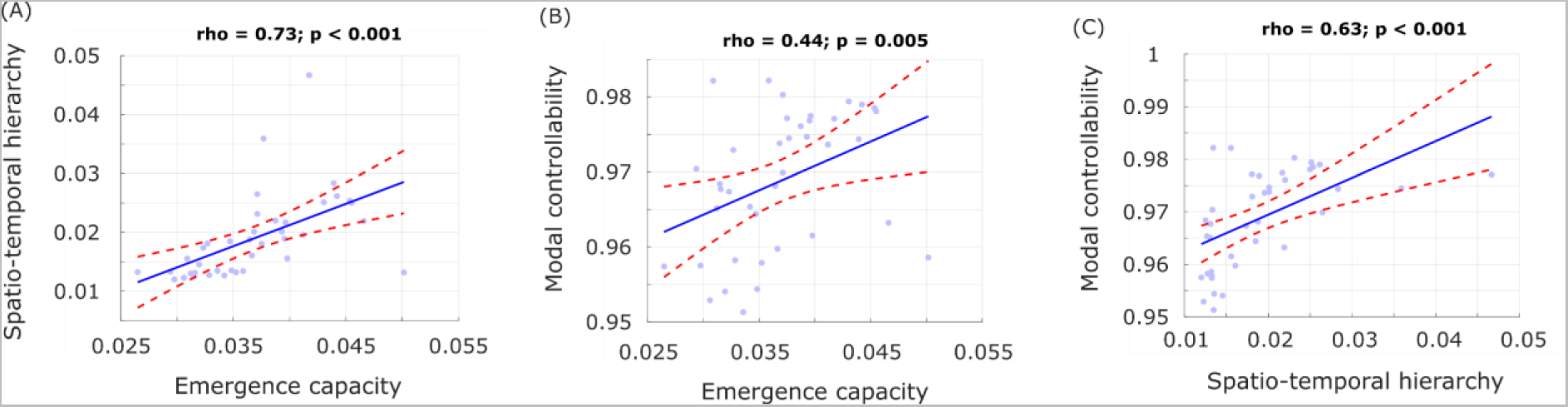
Functional and structural properties of the brain are correlated across subjects. Spearman’s rank-based correlation tests between each pair of structural and functional measures that had exhibited significant differences across DOC patients and controls. Each data-point represents one subject (note that the two healthy controls who did not have functional data were not included in this analysis).

**Figure 5.**
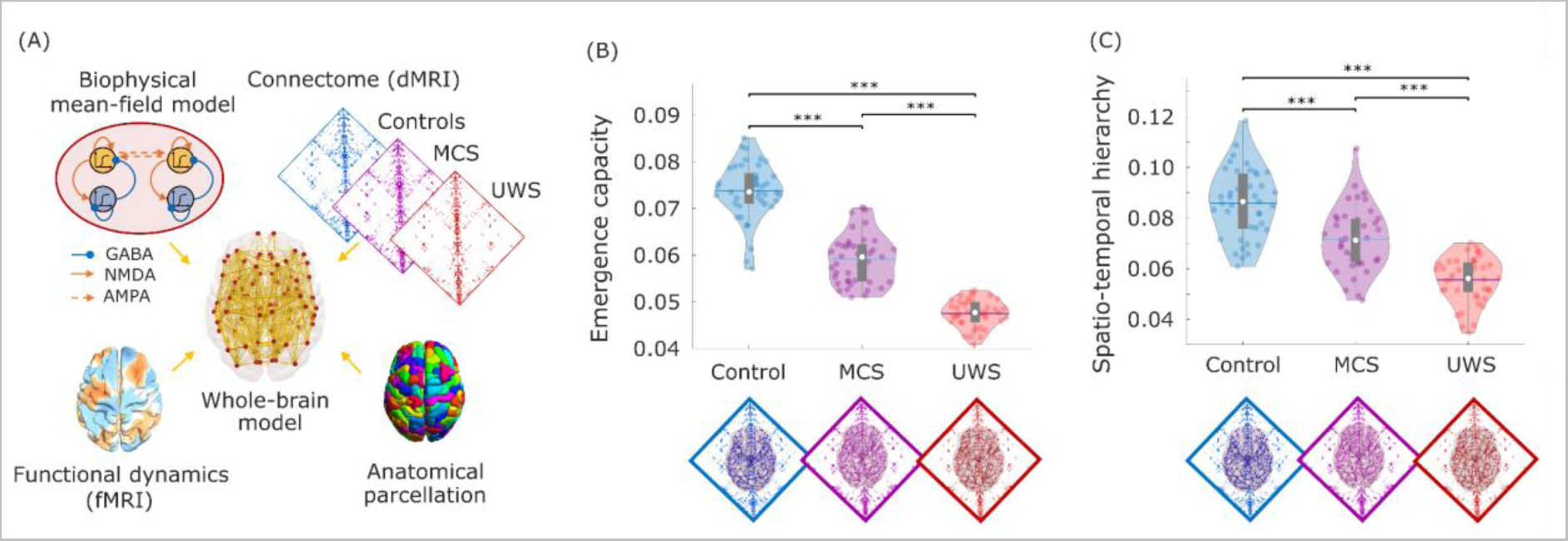
Whole-brain models informed by empirical connectomes replicate empirical changes in brain dynamics. (A) Overview of the whole-brain modelling approach to investigate structure-function relationships. The whole-brain model is based on local biophysical models of excitatory and inhibitory neuronal populations, corresponding to brain regions, interconnected by a network of structural connections obtained from diffusion MRI from each group of subjects (healthy controls, MCS and UWS patients). The whole-brain model has one free parameter, the global coupling *G,* which is selected as the value just before the simulated firing rate becomes unstable. (B) Emergence capacity is highest in the dynamics simulated from control connectome, in line with empirical results. (C) Spatio-temporal hierarchical character is highest in the dynamics simulated from control connectome, in line with empirical results. Each data-point corresponds to one of 40 simulations obtained from each whole-brain model. White circle, median; center line, mean; box limits, upper and lower quartiles; whiskers, 1.5x interquartile range. *** p < 0.001, FDR-corrected.

### Causal evidence for structure-function relationships from whole-brain computational models

Finally, we sought to determine whether the structural alterations observed in DOC patients may be part of the causal mechanism responsible for the observed functional deficits (emergence capacity and spatiotemporal hierarchy). To this end we employed whole-brain computational modelling, a powerful tool to investigate how macroscale neural dynamics emerge from the underlying anatomical connectivity ^15, 38–43^. These models represent regional macroscale activity in terms of two key ingredients: (i) a biophysical model of each region’s local dynamics; and (ii) inter-regional anatomical connectivity. In particular, the neurobiologically plausible Dynamic Mean Field (DMF) model relies on a mean-field reduction to recapitulate the microscale neurophysiological properties of spiking neurons ^64–72^. Each cortical region is modelled as a macroscopic neural field comprising mutually coupled excitatory and inhibitory populations, and regions are then connected according to empirical anatomical connectivity obtained e.g., from diffusion weighted imaging (DWI) data ^64–69^. The flexibility of this neurobiologically inspired whole-brain modelling makes it ideal to investigate how the anatomical connectivity of the brain shapes its macroscale neural dynamics ^38, 42, 73^.

We fitted three whole-brain DMF models, each using a connectome obtained from combining the DTI of healthy controls, MCS patients, and UWS patients, respectively. The DMF model has one free parameter, the global coupling *G*; this parameter was selected separately for each model to lie just before the point where the simulated firing rate becomes unstable, which is typically where the model best reproduces empirical brain dynamics (Methods). Analyses performed on 40 simulations generated by each of these models replicated our main empirical findings, showing significant differences in emergence capacity and spatiotemporal hierarchy across each group - being highest in the model derived from healthy connectomes, and lowest in the model derived from UWS connectomes. We chose this data-agnostic tuning procedure to ensure that the results could be unequivocally attributed solely to the structural connectome. However, analogous results were also obtained when fitting the model *G* parameter to best match the empirical dynamics observed in the corresponding condition (Figure S6). As with the empirical results, the results pertaining to global emergence capacity and spatio-temporal hierarchy could also be replicated using the 129-ROI parcellation (Figure S7). Overall, this computational modelling demonstrates that injury-induced changes in the structural connectome are sufficient to replicate the corresponding alterations in functional brain dynamics observed in chronically unresponsive patients.

## Discussion

The relationship between mind and emergence has been a recurrent open question in the philosophy of mind and cognitive science literature, but heated debates still persist — fostered by the lack of a practical operationalisation of emergence applicable to empirical neuroimaging data ^2^. Here, we present an empirical investigation of this long-standing question in neuroscience. We applied the recent framework of Integrated Information Decomposition to quantify the capacity of macroscale brain activity (from functional MRI recordings) to exhibit emergent phenomena.

Our results reveal that the capacity for causal emergence across the brain is significantly reduced following severe brain injury leading to chronic unresponsiveness. Subsequently, we investigated functional and structural correlates of emergence in the human brain. Functionally, our results show that the brains of chronically unresponsive patients are characterised by diminished hierarchical organisation in the brain’s ability to ignite distributed neural activity. To explore how these functional alterations are related to injury-induced changes in the brain’s structural organisation, we used structural connectivity data obtained from diffusion-weighted MRI to examine differences in controllability of the structural connectome in DOC patients. Our investigation revealed that the organisation of DOC patients’ structural brain networks exhibits a consistent reduction in modal controllability, reflecting diminished structural support to achieve the desired functional configurations. In turn, this reduction in controllability of structural brain networks is associated with the observed functional reductions in emergence capacity and spatiotemporal hierarchy. Finally, a mechanistic relationship between structural and functional changes was confirmed by whole-brain computational modelling, which provided evidence that the kinds of structural alterations observed in chronically unresponsive patients are sufficient to induce the corresponding deficits in functional emergence and hierarchy.

In this context, it is noteworthy that of the two kinds of structural controllability investigated here, DOC patients exhibited global reductions in modal controllability. Modal controllability reflects the capability of a structural network to support transitions to functional configurations that are very different from the current one and therefore difficult to reach ^23^. Whereas transitions between similar configurations may be the result of mere spontaneous fluctuations within each region’s activity, transitions between distant configurations are not likely to arise from such a localised mechanism - instead, their occurrence represents a whole-brain, global phenomenon. In particular, we found that brains that are more modally controllable also exhibit greater hierarchical character (variability across regions) in terms of the capacity for local intrinsic events to ignite global propagation. We speculate that this ability to navigate between distant configurations via non-trivial dynamics may facilitate the presence of causally emergent dynamics - an intriguing possibility that opens several avenues for theoretical and empirical inquiry.

Accordingly, the combination of compromised modal controllability, hierarchical ignition, and emergence suggest that DOC patients’ brain dynamics may suffer from insufficient structural support for transitions towards distant functional configurations. In other words, we speculate that lack of modal controllability at the structural level results in diminished capacity for hierarchical activity and ultimately emergence. This might explain why DOC patients remain chronically unresponsive, unlike anaesthetised or asleep individuals, who also exhibit behavioural unresponsiveness and diminished hierarchical character of ignition, but only temporarily. Sleep and anaesthesia do not influence the connectome, which is therefore still capable in principle of supporting major changes in configuration, and the corresponding capacity for emergent dynamics. In contrast, the results of our whole-brain simulations indicate that the connectomes of DOC patients are less capable of supporting hierarchical and emergent brain dynamics - which seems to be critical for supporting consciousness and higher-order cognition, in line with recent proposals ^74^.

### Limitations and future directions

A number of limitations should be acknowledged when interpreting the results of the present study. Firstly, the account of emergence capacity adopted here is based on Integrated Information Decomposition, which is a recent development in the field of information theory and may be subject to further refinements as this field evolves ^5^. In particular, although we have shown that our results are robust to the use of different operationalisations of information decomposition, methods for estimating ΦID in empirical data are not yet capable of accounting for all brain regions simultaneously, and therefore here we opted to use the average of all pairwise interactions as our quantification of global capacity to support causal emergence. Thus, we acknowledge that an important avenue for future work will be to extend our approach beyond pairwise interactions, and quantify causal emergence across larger groups of regions, up to the entire brain simultaneously, whether through theoretical developments or computational approximations. Indeed, recent advances in the related but complementary account of emergence proposed by Integrated Information Theory ^75–78^ present promising avenues for future investigation. Similarly, “hierarchy” is a protean, multi-faceted concept in neuroscience ^7, 16^, and different operationalisations may bear different relationships with emergence, which should be borne in mind when interpreting the present results.

Additionally, although here we capitalised on the availability of functional and diffusion MRI data in the same cohort of patients, future work may also seek to investigate emergence capacity from electrophysiological signals, which have higher temporal resolution and provide a more direct quantification of neuronal activity. It is also worth acknowledging that the network control framework adopted here for the analysis of structural connectivity rests on assumptions about linearity ^23, 51^. While recent evidence suggests that fMRI data may be adequately captured by linear accounts ^79, 80^, it would also be interesting to consider non-linear extensions of the tools from network control theory employed here.

Finally, it is worth acknowledging that our results did not always identify statistically significant differences between healthy controls and MCS patients, or between MCS and UWS patients. We believe that this is likely due in part to our limited sample sizes and statistical stringency, since the larger sample sizes allowed by whole-brain modelling (40 for each group’s connectome) provided statistically significant differences between each group. Additionally, our correlation plots suggest that patients and controls may lie on a continuum in terms of these functional and structural characteristics of the brain, rather than occupying clearly defined categories. Thus, replication in a larger cohort may be warranted to shed light on the differences between patient subgroups. In this context, future work may investigate whether the structural connectomes of DOC patients who recover consciousness also show a corresponding recovery of modal controllability, and whether emergence from unconsciousness also corresponds to restored emergence in the brain’s functional dynamics.

## Conclusion

Overall, in the present work we combined a suite of cutting-edge computational tools to characterise emergence capacity in the human brain, in the context of functional and structural changes induced by severe brain injury. Bringing the notion of emergence from the realm of philosophy into neuroscience, we identified links between emergence and the spatio-temporal hierarchy of local-global interactions in the human brain, and further discovered a fundamental role of the structural connectome in supporting emergent and hierarchical dynamics. Taken together, the present results lead us to speculate that the chronic nature of unconsciousness in DOC patients may be due to permanent impairment of the fundamental neural infrastructures required to support hierarchical brain dynamics, capable of balancing local segregation and global integration - and ultimately the emergence of consciousness.

## Methods

### Disorders of Consciousness Patient Data

The DOC patient data employed in this study have been published before ^25, 26, 71, 81, 82^. For clarity and consistency of reporting, where applicable we use the same wording as our previous studies.

#### Recruitment

As previously reported ^26^, 71 DOC patients were recruited from specialised long-term care centres from January 2010 to December 2015. Ethical approval for this study was provided by the National Research Ethics Service (National Health Service, UK; LREC reference 99/391). Patients were eligible to be recruited in the study if they had a diagnosis of chronic disorder of consciousness, provided that written informed consent to participation was provided by their legal representative, and provided that the patients could be transported to Addenbrooke’s Hospital (Cambridge, UK). The exclusion criteria included any medical condition that made it unsafe for the patient to participate, according to clinical personnel blinded to the specific aims of the study; or any reason that made a patient unsuitable to enter the MRI scanner environment (e.g., non-MRI-safe implants). Patients were also excluded based on significant pre-existing mental health problems, or insufficient fluency in the English language prior to their injury. After admission to Addenbrooke’s Hospital, each patient underwent clinical and neuroimaging testing, spending a total of five days in the hospital (including arrival and departure days). Neuroimaging scanning took place at the Wolfson Brain Imaging Centre (Addenbrooke’s Hospital, Cambridge, UK), and medication prescribed to each patient was maintained during scanning.

For each day of admission, Coma Recovery Scale-Revised (CRS-R) assessments were recorded at least daily. Patients whose behavioural responses were not indicative of awareness at any time, were classified as UWS. In contrast, patients were classified as being in a minimally conscious state (MCS) if they provided behavioural evidence of simple automatic motor reactions (e.g., scratching, pulling the bed sheet), visual fixation and pursuit, or localisation to noxious stimulation. Since this study focused on whole-brain properties, coverage of most of the brain was required, and we followed the same criteria as in our previous studies ^25, 26, 71^: before analysis took place, patients were systematically excluded if an expert neuroanatomist blinded to diagnosis judged that they displayed excessive focal brain damage (over one third of one hemisphere), or if brain damage led to suboptimal segmentation and normalisation, or due to excessive head motion in the MRI scanner (exceeding 3mm translation or 3 degrees rotation). One additional patient was excluded due to incomplete acquisition. A total of 21 adults (13 males; 17-70 years; mean time post injury: 13 months) meeting diagnostic criteria for unresponsive wakefulness syndrome/vegetative state (UWS; N = 10) or minimally conscious state (MCS; N = 11) due to brain injury were included in this study (Table 1).

**Table 1:** Demographic information for patients with Disorders of Consciousness. CRS-R, Coma Recovery Scale-Revised; UWS, Unresponsive Wakefulness Syndrome; MCS, Minimally Conscious State; TBI, Traumatic Brain Injury.

| Sex | Age | Aetiology | Diagnosis | CRS-R<br>Score | Scan |
| --- | --- | --- | --- | --- | --- |
| M | 46 | TBI | UWS | 6 | 12 dir |
| M | 57 | TBI | MCS | 12 | 12 dir |
| M | 35 | Anoxic | UWS | 8 | 12 dir |
| M | 17 | Anoxic | UWS | 8 | 12 dir |
| F | 31 | Anoxic | MCS | 10 | 12 dir |
| F | 38 | TBI | MCS | 11 | 12 dir |
| M | 29 | TBI | MCS | 10 | 63 dir |
| M | 23 | TBI | MCS | 7 | 63 dir |
| F | 70 | Cerebral bleed | MCS | 9 | 63 dir |
| F | 30 | Anoxic | MCS | 9 | 63 dir |
| F | 36 | Anoxic | UWS | 8 | 63 dir |
| M | 22 | Anoxic | UWS | 7 | 63 dir |
| M | 40 | Anoxic | UWS | 7 | 63 dir |
| F | 62 | Anoxic | UWS | 7 | 63 dir |
| M | 46 | Anoxic | UWS | 5 | 63 dir |
| M | 21 | TBI | MCS | 11 | 63 dir |
| M | 67 | TBI | MCS | 11 | 63 dir |
| F | 55 | Hypoxia | UWS | 7 | 63 dir |
| M | 28 | TBI | MCS | 8 | 63 dir |
| M | 22 | TBI | MCS | 10 | 63 dir |
| F | 28 | ADEM | UWS | 6 | 63 dir |

#### FMRI Data Acquisition

As previously reported ^25, 26, 71^, resting-state fMRI was acquired for 10 minutes (300 volumes, TR=2000ms) using a Siemens Trio 3T scanner (Erlangen, Germany). Functional images (32 slices) were acquired using an echo planar sequence, with the following parameters: 3 x 3 x 3.75mm resolution, TR = 2000ms, TE = 30ms, 78 degrees FA. Anatomical scanning was also performed, acquiring high-resolution T1-weighted images with an MPRAGE sequence, using the following parameters: TR = 2300ms, TE = 2.47ms, 150 slices, resolution 1 x 1 x 1mm.

#### Acquisition of Diffusion-Weighted Imaging Data

As we previously reported ^25, 71^, the DOC patients’ data were acquired over the course of several years, and as a result two different diffusion-weighted image acquisition schemes were used. The first acquisition scheme involved diffusion-sensitising gradients applied along 12 non-collinear directions, and 5 different b-values ranging from 340 to 1590 s/mm^2^. An echo planar sequence was used (TR = 8300 ms, TE = 98 ms, matrix size = 96 x 96, 63 slices, slice thickness = 2 mm, no gap, flip angle = 90°). This acquisition scheme was used for the first N=6 patients (Table 1). The second acquisition scheme included 63 directions with a b-value of 1000 s/mm2; this acquisition scheme was adopted for all remaining DOC patients and also for all healthy controls. Each of these DWI acquisition types has been used before with DOC patients ^25, 34, 35, 71^.

### Healthy Controls

We also used previously-acquired fMRI and DWI data from N=20 healthy volunteers (13 males; 19-57 years), with no history of psychiatric or neurological disorders ^25^. The mean age was not significantly different between healthy controls (M = 35.75; SD = 11.42) and DOC patients (M = 38.24; SD = 15.96) (*t*(39) = −0.57, *p* = 0.571, Hedges’s *g* = −0.18; permutation-based t-test).

#### FMRI Data Acquisition

Resting-state fMRI was acquired for 5:20 minutes (160 volumes, TR=2000ms) using a Siemens Trio 3T scanner (Erlangen, Germany). The acquisition parameters were the same as those for the DOC patients: Functional images (32 slices) were acquired using an echo planar sequence, with the following parameters: 3 x 3 x 3.75mm resolution, TR = 2000ms, TE = 30ms, 78 degrees FA. High-resolution T1-weighted anatomical images were also acquired, using an MPRAGE sequence with the following parameters: TR = 2300ms, TE = 2.47ms, 150 slices, resolution 1 x 1 x 1mm. Data from two subjects were excluded due to incomplete acquisition, leaving N=18 healthy controls for the functional analysis.

#### Acquisition of Diffusion-Weighted Imaging Data

The diffusion-weighted acquisition scheme was the same 63-directions scheme used for the DOC patients, as described above and in previous work (Luppi et al., 2021): TR = 8300 ms, TE = 98 ms, matrix size = 96 x 96, 63 slices, slice thickness = 2 mm, no gap, flip angle = 90°, 63 directions with a b-value of 1000 s/mm2.

### Data preprocessing and denoising

#### Functional MRI data

We preprocessed the functional imaging data using the CONN toolbox, version 17f (http://www.nitrc.org/projects/conn) ^83^ based on Statistical Parametric Mapping 12 (http://www.fil.ion.ucl.ac.uk/spm). For each dataset and condition, we applied a standard preprocessing pipeline, the same as we employed in our previous studies ^26, 62, 84, 85^. The pipeline involved the following steps: removal of the first five volumes, to achieve steady-state magnetization; motion correction; slice-timing correction; identification of outlier volumes for subsequent scrubbing by means of the quality assurance/artifact rejection software *art* (http://www.nitrc.org/projects/artifact_detect); normalisation to Montreal Neurological Institute (MNI-152) standard space (2 mm isotropic resampling resolution), using the segmented grey matter image from each volunteer’s T1-weighted anatomical image, together with an *a priori* grey matter template. Due to the presence of deformations caused by brain injury in the DOC patients, our preprocessing avoided automated pipelines. Each patient’s brain was individually preprocessed using SPM12, with visual inspections after each step. Additionally, to further reduce potential movement artefacts, data underwent despiking with a hyperbolic tangent squashing function. These procedures are the same as in our previous publications on these data ^26, 62, 71^.

To reduce noise due to cardiac and motion artefacts, we applied the anatomical CompCor method of denoising the functional data ^86^. The anatomical CompCor method (also implemented within the CONN toolbox) involves regressing out of the functional data the following confounding effects: the first five principal components attributable to each individual’s white matter signal, and the first five components attributable to individual cerebrospinal fluid (CSF) signal; six subject-specific realignment parameters (three translations and three rotations) as well as their first-order temporal derivatives; the artefacts identified by *art;* and main effect of scanning condition. Linear detrending was also applied, and the subject-specific denoised BOLD signal timeseries were band-pass filtered to eliminate both low-frequency drift effects and high-frequency noise, thus retaining frequencies between 0.008 and 0.09 Hz. The step of global signal regression (GSR) has received substantial attention in the literature as a denoising method ^87–89^. GSR mathematically mandates that approximately 50% of correlations between regions will be negative ^90^; however, the proportion of anticorrelations between brain regions has been shown to vary across states of consciousness, including anaesthesia and DOC ^25, 71, 91^. Indeed, recent work has demonstrated that the global signal contains information about pathological and pharmacological states of unconsciousness ^92^. Therefore, in line with our previous studies, here we decided to avoid GSR in favour of the aCompCor denoising procedure, which is among those recommended for investigations of brain dynamics ^88^.

#### DWI Preprocessing and Tractography

The diffusion data were preprocessed with MRtrix3 tools, following the same pipeline as in our previous work ^25, 71, 91^. After manually removing diffusion-weighted volumes with substantial distortion ^35^, the pipeline involved the following steps: (i) DWI data denoising by exploiting data redundancy in the PCA domain ^93^ (*dwidenoise* command); (ii) correction for distortions induced by eddy currents and subject motion by registering all DWIs to b0, using FSL’s *eddy* tool (through MRtrix3 *dwipreproc* command); (iii) rotation of the diffusion gradient vectors to account for subject motion estimated by *eddy* ^94^; (iv) b1 field inhomogeneity correction for DWI volumes (*dwibiascorrect* command); and (v) generation of a brain mask through a combination of MRtrix3 *dwi2mask* and FSL *BET* commands.

After preprocessing, the DTI data were reconstructed using the model-free q-space diffeomorphic reconstruction algorithm (QSDR) implemented in DSI Studio (www.dsi-studio.labsolver.org) ^95^, following our previous work ^25, 71, 96^. Use of QSDR is desirable when investigating group differences ^95, 97, 98^ because this algorithm preserves the continuity of fiber geometry for subsequent tracking ^95^, since it reconstructs the distribution of the density of diffusing water in standard space. This approach has therefore been adopted in previous connectomics studies focusing on healthy individuals ^23^ but also brain-injured patients ^99^and DOC patients specifically ^95, 97, 98^. QSDR initially reconstructs DWI data in native space, and subsequently computes values of quantitative anisotropy (QA) in each voxel, based on which DSI Studio performs a nonlinear warp from native space to a template QA volume in Montreal Neurological Institute (MNI) space. Once in MNI standard space, spin density functions are reconstructed, with a mean diffusion distance of 1.25 mm with three fiber orientations per voxel^95^.

Finally, fiber tracking was carried out by means of DSI Studio’s own “FACT” deterministic tractography algorithm, requesting 1,000,000 streamlines according to widely adopted parameters ^23, 25, 71, 99^: angular cutoff = 55◦, step size = 1.0 mm, tract length between 10mm (minimum) and 400mm (maximum), no spin density function smoothing, and QA threshold determined by DWI signal in the cerebro-spinal fluid. Streamlines were automatically rejected if they presented improper termination locations, based on a white matter mask automatically generated by applying a default anisotropy threshold of 0.6 Otsu’s threshold to the anisotropy values of the spin density function ^23, 25, 71, 99^.

### Brain parcellation

For both BOLD and DWI data, brains were parcellated into 234 cortical and subcortical regions of interest (ROIs), according to the Lausanne sub-parcellation of the Desikan-Killiany anatomical atlas ^59,100^. This parcellation has been used in previous work on controllability of structural brain networks ^23, 49^. Recent work has shown that parcellations in the range of 200 regions provide generalisable network results ^96^. Additionally, to ensure the robustness of our results, we also replicated our analyses with a different version of the same parcellation, which includes 129 cortical and subcortical regions ^59^.

### Quantifying emergence capacity

Consider a stochastic process 𝑋 comprised of two random variables evolving jointly over time, 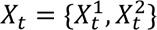. In our case, this corresponds to the timeseries of the BOLD activity of two brain regions, although in other applications it could be any form of multivariate timeseries data. One can now consider the amount of information flowing from the system’s past to its future, known as time-delayed mutual information (TDMI) and given by 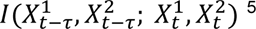.

Following the insights of Williams and Beer ^101^, the information that two source variables *X* and *Y* give about a third target variable *Z*, denoted by *I*(*X,Y* ; *Z*), can be decomposed in terms of different *types* of information: information provided by one source but not the other (unique information), by both sources separately (redundant information), or jointly by their combination (synergistic information). The mathematical framework of Integrated Information Decomposition (ΦID) ^5^ has generalised this insight to the case of multiple sources and multiple target variables - such as the respective future states of the parts of the system under consideration. Thus, through ΦID it is possible to decompose TDMI into redundant, unique, and synergistic information shared with respect to both past and present state of both variables.

Importantly, ΦID’s decomposition of information offers a way to compute the formal, quantitative definition of causal emergence established by Rosas and colleagues ^1^, according to which a supervenient feature 𝑉_𝑡_ of system *X* is *causally emergent* if it has predictive power about the future evolution of 𝑋_𝑡_ that is unique with respect to 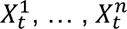. Here, *supervenience* of 𝑉_𝑡_ on 𝑋_𝑡_ (the instantaneous state of the system at time *t*) is defined as 𝑉_𝑡_ being a function of 𝑋_𝑡_, so that there is nothing about 𝑉_𝑡_ that can be predicted from the system’s previous state, 𝑋_𝑡−1_, that cannot be already predicted from the system’s current state, 𝑋_𝑡_ ^1^. Crucially, it can be mathematically demonstrated ^1^ that a system’s capacity for having causally emergent features depends directly on how synergistic its dynamics are. In particular, from ΦID’s characterisation of causal emergence it follows that causal emergence can take place in two distinct scenarios: when an emergent feature has unique predictive power over parts of the system (“*downward causation*”), or when the emergent feature’s unique predictive power is not over any individual constituent but only over the system as a whole (“*causal decoupling*”). The latter can be thought of as “the macroscale having causal influence on the macroscale, above and beyond the microscale effects” ^1^. Here we focus on the system’s “emergence capacity”, the combination of both downward causation and causal decoupling.

In practice, our method for computing the emergence capacity involves obtaining the full integrated information decomposition of the system, which is achieved by setting up a linear system of 15 equations with 16 unknowns relating various standard (Shannon) mutual information terms with the redundant, unique, and synergistic components of the TDMI. The system can be solved by specifying the redundancy between 𝑋_𝑡−𝜏_ and 𝑋_𝑡_ which we compute following the “common change in surprisal” (CCS) method ^102^. This allows us to solve the linear system of equations and obtain all components of the Integrated Information Decomposition of our system of interest, from which we obtain emergence capacity as the sum of its constituent atoms (downward causation + causal decoupling). For all the analyses in the paper we use the JIDT toolbox ^103^ to compute all information-theoretic quantities for each pair of brain regions, using a standard plug-in estimator applied to the mean-binarised BOLD signals. To validate our results, we also replicated them using continuous instead of discrete signals and the Gaussian solver implemented in JIDT ^103^. Likewise, we replicated our results using an alternative definition of redundancy known as the minimum mutual information (MMI) ^101^. In accordance with our previous work ^6, 104^ and previous studies using information-theoretic measures in the context of functional MRI data, for these analyses we used a state-of-the-art toolbox ^105^ to deconvolve the hemodynamic response function from our regional BOLD signal timeseries.

### Spatiotemporal Hierarchy from Intrinsic-driven Ignition

“Intrinsic-driven ignition” ^14^ quantifies the extent to which spontaneously occurring (“intrinsic”) local events elicit whole-brain activation (“ignition”). For this analysis, first the BOLD signal is narrowband-filtered in the range 0.04-0.07 Hz range, in line with previous ^1^. The filtered timeseries are then transformed into z-scores, and subsequently thresholded to obtain a binary sequence σ based on the combined mean and standard deviation of the regional transformed signal, such that σ(t) = 1 if z(t) > 1 and is crossing the threshold from below, indicating that a local event has been triggered; otherwise, σ(t) = 0 ^1^. Note that the threshold of 1 standard deviation for triggering an event is chosen for consistency with previous work, but it has been demonstrated that the results of this procedure are robust to the specific threshold chosen ^106^. Subsequently, for each brain region, when that region triggers a local event (σ(t) = 1, (“driver event” in Figure 2A), the resulting global ignition is computed within a time-window of 4 TRs (corresponding approximately to the duration of one hemodynamic response function, given our TR of 2s). An NxN binary matrix M is then constructed (Figure 2A), indicating whether in the period of time under consideration two regions *i* and *j* both triggered an event (M*_ij_ =* 1). The size of the largest connected component of this binary matrix M defines the breadth of the global ignition generated by the driver region at time *t,* termed “intrinsic-driven ignition” (IDI) ^14^ (Figure 2A). To obtain a measure of spatio-temporal hierarchy of local-global integration, each region’s IDI values are averaged over time, and the variability (standard deviation) across regions is then computed. Consequently, higher standard deviation reflects more heterogeneity across brain regions with respect to their capability to induce ignition, which suggests in turn a more elaborate hierarchical organisation between them (Figure 2A).

### Structural network construction

A network consists of two basic elements: nodes, and the edges connecting them. To construct structural brain networks, patients’ brains were parcellated into 234 or 129 cortical and subcortical regions of interest (ROIs) of approximately equal size, derived from the Lausanne atlas (the parcels were dilated by 2 voxels to extend them to the grey-matter-white matter interface) ^23^. These ROIs represent the nodes of the brain network. Then, for each pair of nodes *i* and *j*, an edge was drawn between them if there were white matter tracts connecting the corresponding brain regions end-to-end; edge weights were quantified as the number of streamlines connecting each pair of regions end-to-end. In turn, this network can be represented as an adjacency matrix **A**, whose entry Aij corresponds to the weight of connection between nodes (brain regions) *i* and *j*.

### Network Controllability

The model of brain dynamics used for network controllability analysis is based on extensive prior work demonstrating its wide applicability in health and disease ^23, 47–55^. In effect, there exists substantial evidence that linear models provide an adequate description of the brain dynamics measured with fMRI - such that more complicated non-linear models only capture little additional variance ^79, 80^. Additionally, the controllability framework adopted here has been shown to have substantial overlap with the analysis of systems of non-linear oscillators connected with neurobiologically realistic coupling constants (using white matter connectivity, analogous to the use of white matter connectivity employed here) ^107^. Based on this literature and the well-known tractability of linear models, here we follow prior work on network control theory applications to structural brain networks ^23, 99^ which uses a time-invariant network model on discrete-time of the form

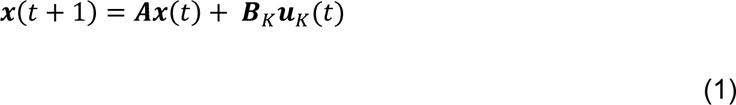

which has been previously used as a model of both BOLD signals and neural activity ^23, 108^. Here, **x** is a vector describing the state of each brain region at a given point in time (e.g. in terms of neural activation as given by the BOLD signal magnitude - though note that the network control framework is agnostic about the nature of the system’s activity), and **A** is the adjacency matrix representing the structural connectome (to ensure Schur stability, the adjacency matrix is divided by its largest singular value + 1 ^23, 99^). In turn, the input **u**_k_ represents the control strategy, which is applied according to the control points K identified by the matrix **B**_K_, where *K* = {*k_1_…k_m_*} and **B**_K_ = [𝒆_𝑘_ , … 𝒆_𝑘_ ] with *e_i_* representing the *i*^th^ canonical vector of size N. While this is a time-discrete model, previous work has shown that the controllability Gramian (see below) is statistically similar to that obtained from continuous-time system ^23, 99^.

Network control analysis enables us to investigate the ability of each brain region to influence the brain’s dynamics in different ways. Technically, the “controllability” of a dynamical system (such as the human brain) refers to the extent to which the state of the dynamical system in question can be driven towards a chosen target state by means of an external input. Based on well-known results from control theory, the system described in Eq. (1) is controllable from the control nodes *K* if the “controllability Gramian” matrix given by

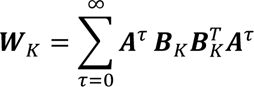

is invertible. Following previous work, the input nodes are chosen one at a time, so that the input matrix **B**_K_ reduces to a vector denoting the control node.

Based on this controllability framework, we focus on two complementary control strategies for determining how the system can be moved towards different states (i.e., regional activation patterns): “average” and “modal” controllability ^23^.

#### Average controllability

If the states that are accessible to the system are conceptualised as constituting an energy landscape, then average controllability describes how easily the system can transition between nearby states on this landscape. Average controllability of a network then equals to the average input energy needed at a set of control nodes, averaged over all possible target states. It is well-established that average input energy is proportional to Trace 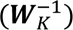. However, since the trace of the inverse Gramian is often uncomputable due to ill-conditioning, we follow previous work ^23^ in using Trace(𝑾_𝐾_) instead (which encodes the energy of the network impulse

response), since the traces of the Gramian and its inverse are inversely proportional.

#### Modal controllability

Modal controllability describes how easily the system can be induced to transition to a state that is distant on the energy landscape of its possible state. Technically, it corresponds to the ability of a node to control each of the dynamic modes of the network, and it can be computed from the matrix **V** of the eigenvectors of **A**. From well-established results, it is known that if the entry *v_ij_* is small, then the *j^th^* mode of the system is poorly controllable from node *i* ^23^. Therefore, here we follow previous work in defining a scaled measure of the controllability from brain region *i* of all the N modes of the system, 𝜆_1_(𝐴) … 𝜆_𝑁_(𝐴) as:

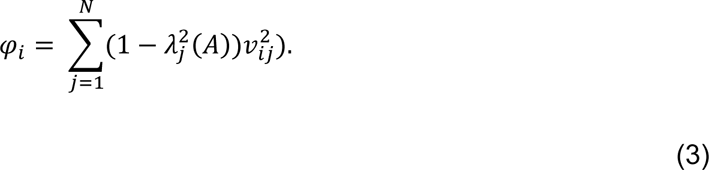

From this definition, a region will have high modal controllability if it is able to control all the dynamic modes of the system, which implies that they are well-suited to drive the system towards difficult-to-reach configurations in the energy landscape ^23^.

For both average and modal controllability, whole-brain values can be obtained by taking the mean of all regional controllability values, as per prior work ^52^.

### Whole-brain computational modelling

Macroscale whole-brain computational models represent regional activity in terms of two key ingredients: (i) a biophysical model of each region’s local dynamics; and (ii) inter-regional anatomical connectivity. Thus, such *in silico* models provide a well-suited tool to investigate how the structural connectivity of the brain shapes the corresponding macroscale neural dynamics ^15, 38–43^. In particular, the Dynamic Mean Field (DMF) model employed here simulates each region (defined via an anatomical parcellation scheme) as a macroscopic neural field comprising mutually coupled excitatory and inhibitory populations (80% excitatory and 20% inhibitory), providing a neurobiologically plausible account of regional neuronal firing rate. Regions are then connected according to empirical anatomical connectivity obtained e.g. from DWI data ^64–66^. The reader is referred to ^68, 70–72^ for details of the DMF model and its implementation. Due to its multi-platform compatibility, low memory usage, and high speed, we used the recently developed *FastDMF* library ^72^, available online at https://www.gitlab.com/concog/fastdmf.

The structural connectivity (SC) for the DMF model used here was obtained by following the procedure described by Wang *et al.* ^43^ to derive a consensus structural connectivity matrix. A consensus matrix ***A*** was obtained separately for each group (healthy controls, MCS patients, UWS patients) as follows: for each pair of regions *i* and *j*, if more than half of subjects had non-zero connection *i* and *j*, ***A****_ij_* was set to the average across all subjects with non-zero connections between *i* and *j*. Otherwise, ***A****_ij_* was set to zero.

The DMF model has one free parameter, known as “global coupling” and denoted by *G*, which accounts for differences in transmission between brain regions, considering the effects of neurotransmission but also synaptic plasticity mechanisms. Thus, separately for each group, we used a model informed by that group’s consensus connectome to generate 40 simulations for each value of *G* between 0.1 and 2.5, using increments of 0.1. Finally, we set the *G* parameter to the value just before the one at which the simulated firing of each model became unstable, reflecting a near-critical regime.

Subsequently, for each group, 40 further simulations were obtained from the corresponding DMF model with the optimal *G* parameter. A Balloon-Windkessel hemodynamic model ^109^ was then used to turn simulated regional neuronal activity into simulated regional BOLD signal. Finally, simulated regional BOLD signal was bandpass filtered in the same range as the empirical data (0.008-0.09 Hz, or 0.04-0.07 Hz for the intrinsic ignition analysis).

As an alternative way of finding the most suitable value of *G* for the simulation of each condition, we adopted the approach previously described ^67, 68, 70, 71^ which aims to obtain the best match between empirical and simulated functional connectivity dynamics. First, we quantified empirical functional connectivity dynamics (FCD) in terms of Pearson correlation between regional BOLD timeseries, computed within a sliding window of 30 TRs with increments of 3 TRs ^67, 68, 70, 71^. Subsequently, the resulting matrices of functional connectivity at times *t_x_* and *t_y_* were themselves correlated, for each pair of timepoints *t_x_* and *t_y_*, thereby obtaining an FCD matrix of time-versus-time correlations. Thus, each entry in the FCD matrix represents the similarity between functional connectivity patterns at different points in time. This procedure was repeated for each subject of each group (controls, MCS, and UWS). For each simulation at each value of *G*, we used the Kolmogorov-Smirnov distance to compare the histograms of empirical (group-wise) and simulated FCD values (obtained from the upper triangular FCD matrix). Finally, we set the model’s *G* parameter to the value that was observed to minimise the mean KS distance - corresponding to the model that is best capable of simulating the temporal dynamics of resting-state brain functional connectivity observed in the corresponding group (Figure S6). After having found the value of *G* for each condition, simulated BOLD signals were obtained as described above.

### Statistical analysis

Statistical significance of differences in functional measures was assessed by conducting a three-way analysis of variance (ANOVA), testing for the effect of interest (diagnostic condition, with three levels: control, MCS and UWS). Upon finding the effect of interest to be statistically significant, we conducted post-hoc tests (two-sided non-parametric between-subjects t-tests with 10,000 permutations) using three pairwise comparisons between the conditions (control vs. MCS, control vs. UWS, and MCS vs. UWS). We adopted the method of Benjamini and Hochberg ^44^ to control the false discovery rate across these three pairwise comparisons, at a two-sided alpha value of 0.05. The effect sizes were estimated using Cohen’s *d*. For the statistical analysis of differences in structural measures (global average and modal controllability), we used an analysis of covariance to control for DWI sequence type (12 vs 63 directions) and the number of removed volumes due to motion corruption, as covariates of no interest ^25^. Although we thoroughly preprocessed our functional data to minimise the potential confounding effects of head motion, to ensure the robustness of our results we also carried out a validation analysis including motion (mean framewise displacement) as a covariate of no interest in the functional analyses.

## Acknowledgements

The authors would like to thank all the participants, the patients, and their families for their contribution to this study. This work was supported by the Gates Cambridge Trust (to AIL), the National Institute for Health Research (NIHR, UK), Cambridge Biomedical Research Centre and NIHR Senior Investigator Awards [to DKM], the Stephen Erskine Fellowship (Queens’ College, Cambridge, to EAS), the Canadian Institute for Advanced Research (CIFAR; grant RCZB/072 RG93193) (to DKM and EAS); the British Oxygen Professorship of the Royal College of Anaesthetists [to DKM] and the Vice-Chancellor Award (to PC). DKM is a Fellow of the CIFAR Brain, Mind, and Consciousness Programme. PAMM and DB are funded by the Wellcome Trust (grant no. 210920/Z/18/Z). FER is funded by the Ad Astra Chandaria foundation. Computing infrastructure at the Wolfson Brain Imaging Centre (WBIC-HPHI) was funded by the MRC research infrastructure award (MR/M009041/1). The research was also supported by the NIHR Brain Injury Healthcare Technology Co-operative based at Cambridge University Hospitals NHS Foundation Trust and University of Cambridge. The views expressed are those of the authors and not necessarily those of the NIHR or the Department of Health and Social Care.

## Author Contributions

AIL: conceived the study; analysed data; wrote first draft of the manuscript. PAM: conceived the study; contributed to data analysis and interpretation of results; reviewed and edited the manuscript. FR: conceived the study; contributed to data analysis and interpretation of results; reviewed and edited the manuscript. M.M.C., P.C., A.R.D.P: contributed to data analysis. DKM: reviewed the manuscript. DB: reviewed and edited the manuscript. EAS: conceived the study; reviewed and edited the manuscript. P.F., G.B.W., J.A., J.D.P., D.K.M. and E.A.S. were involved in designing the original studies for which the present data were collected. P.F., M.M.C., G.B.W., J.A., and E.A.S. all participated in data collection.

