## Supplementary Information for "Reduced emergent character of neural dynamics in patients with a disrupted connectome"

### Supplementary Figures

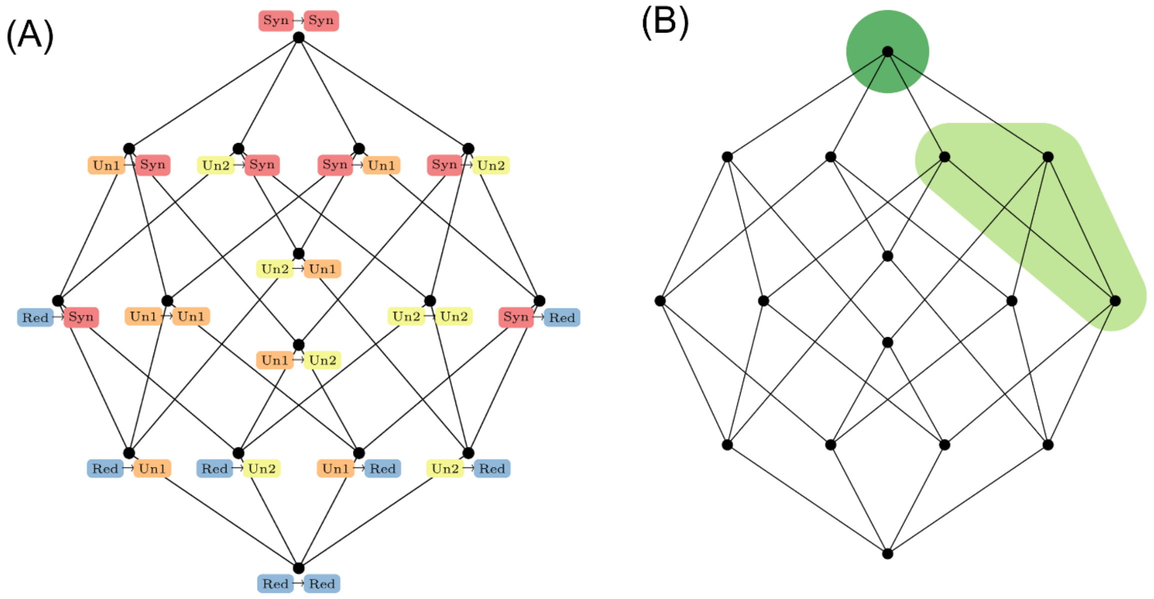


**Figure S1**. Schematic of ΦID's account of causal emergence. (A). 16 "information atoms" representing the possible combinations of synergistic (red), unique (orange and yellow), and redundant (blue) information across time. (B) Information atoms that constitute causal emergence: causal decoupling (dark green) and downward causation (light green).

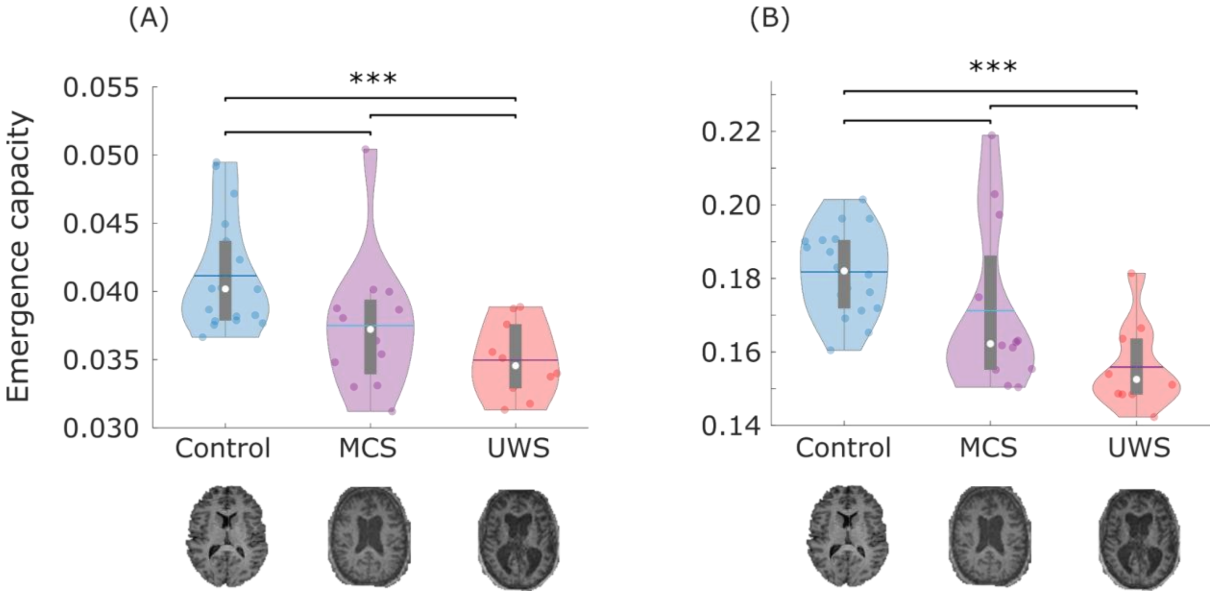


**Figure S2. Replication of emergence results with alternative implementations of information decomposition.** (A) Significant differences in global emergence capacity are observed when performing integrated information decomposition using continuous signals (using the JIDT Gaussian solver). (B) Significant differences in global emergence capacity are observed when performing integrated information decomposition using the minimum mutual information (MMI) definition of redundancy. Data points represent subjects. White circle, median; center line, mean; box limits, upper and lower quartiles; whiskers, 1.5x interquartile range. ** $p < 0.01$; *** $p < 0.001$, FDR-corrected.

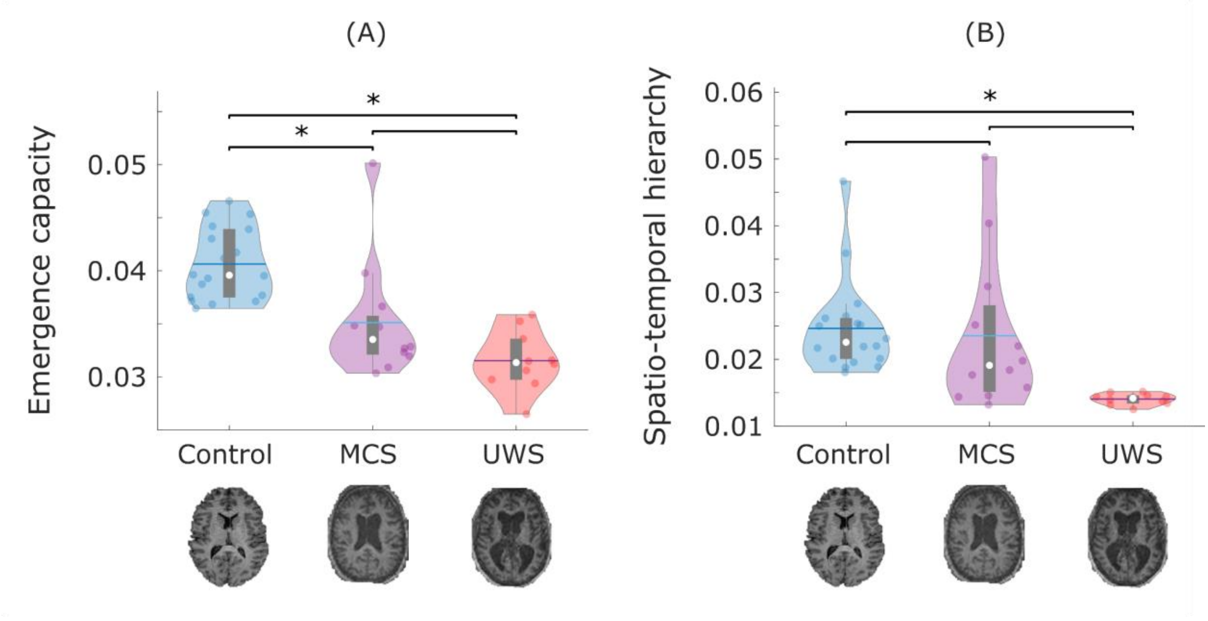


**Figure S3. Functional results after controlling for head motion.** (A) The global emergence capacity. (B) Spatio-temporal hierarchy of intrinsic-driven ignition. Data points represent subjects. White circle, median; center line, mean; box limits, upper and lower quartiles; whiskers, 1.5x interquartile range. ** $p < 0.01$; *** $p < 0.001$, FDR-corrected.

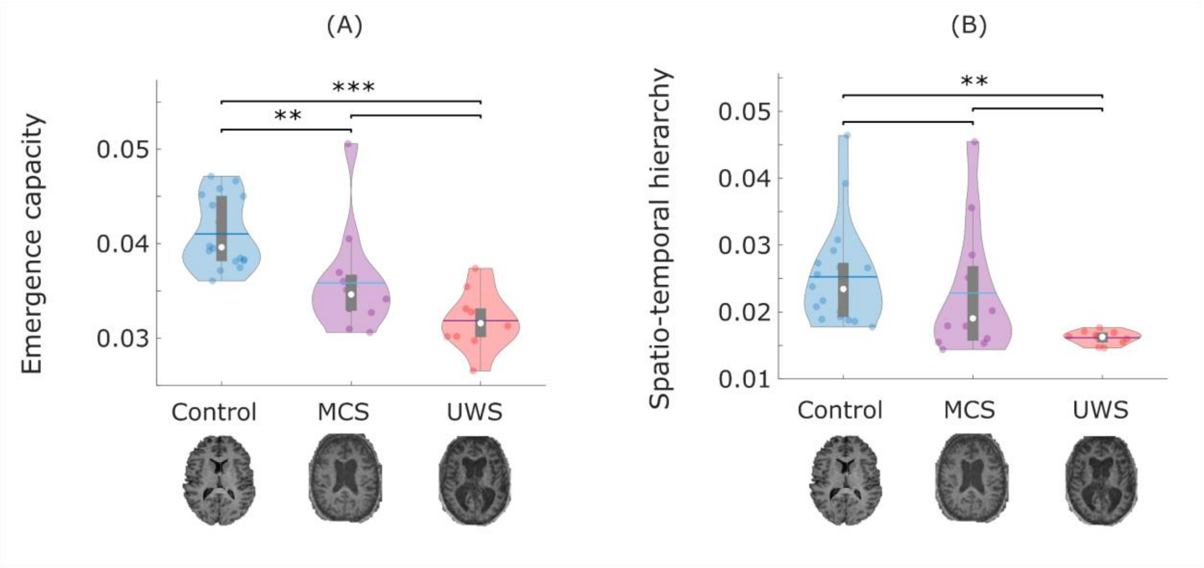


**Figure S4. Replication of empirical results with Lausanne-129 parcellation.** (A) Empirical global emergence capacity**.** (B) Empirical spatio-temporal hierarchy of intrinsic ignition. Data points represent subjects. White circle, median; center line, mean; box limits, upper and lower quartiles; whiskers, 1.5x interquartile range. ** p < 0.01; *** p < 0.001, FDR-corrected.

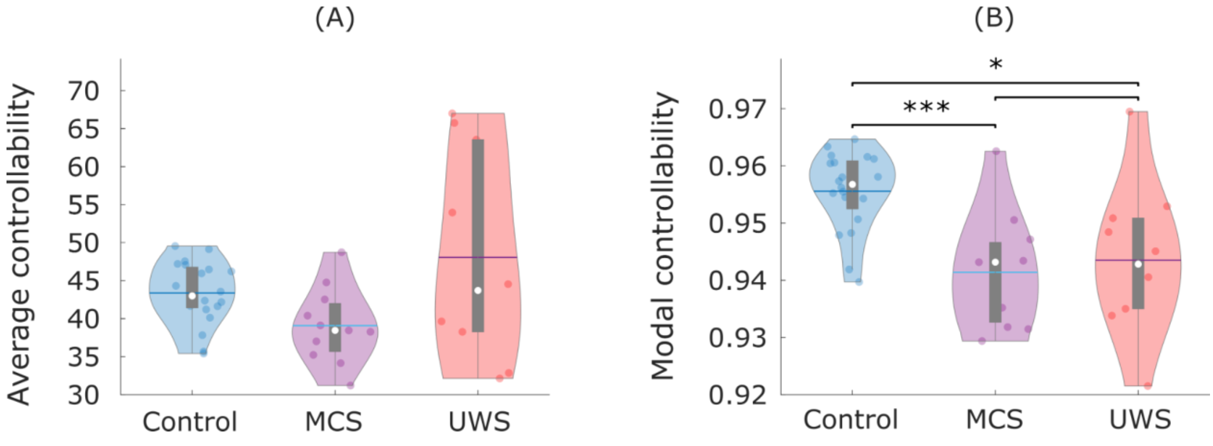


**Figure S5. Replication of structural controllability results with Lausanne-129 parcellation.** (A) No significant effect of group membership (control, MCS, UWS) on global average controllability ($F(2,36) = 1.30$, $p = 0.284$). (B) Global modal controllability is significantly reduced in DOC patients. Data points represent subjects. White circle, median; center line, mean; box limits, upper and lower quartiles; whiskers, 1.5x interquartile range. *** p < 0.001, FDR-corrected.

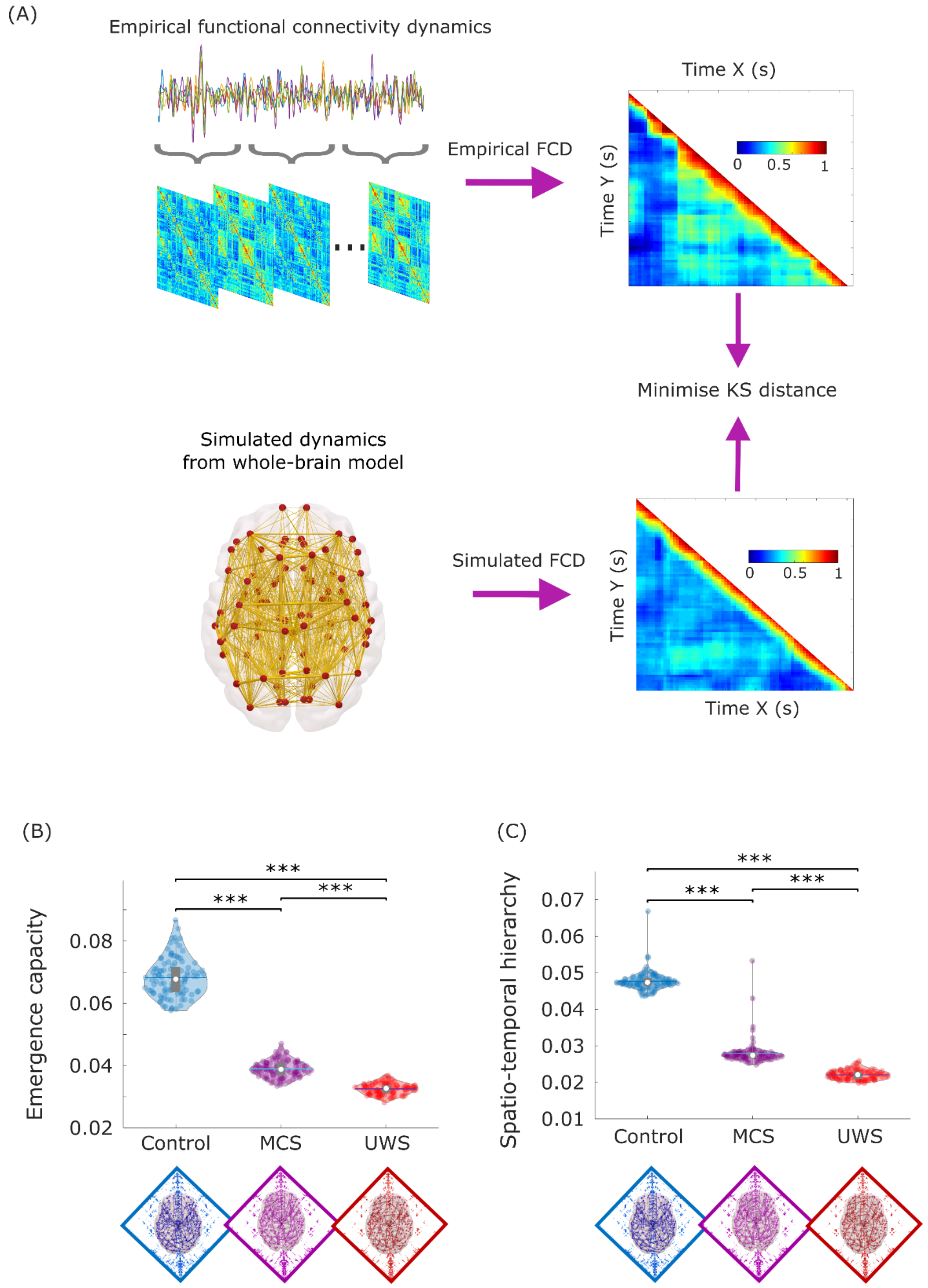


**Figure S6. Simulated results from alternative model fitting procedure.** (A) Overview of model-fitting based on functional connectivity dynamics (FCD). Time-resolved matrices of functional connectivity are obtained from empirical functional MRI via the sliding-window approach: regional BOLD time-series are partitioned into windows of 30 TRs, sliding by 3 TRs at a time, following the same approach as previous work using the DMF model; functional connectivity between each pair of regions is computed within each window by means of Pearson correlation, generating a stack of FC matrices representing the evolution of FC over time. The same procedure is repeated for the simulated BOLD timeseries produced by the model with various levels of the global coupling parameter, *G*. For both the empirical and simulated functional connectivity dynamics (FCD), a time-versus-time FCD matrix is computed by correlating the time-dependent FC matrices centred at each timepoint. Across values of the global coupling parameter *G*, we compute the KS-distance between the empirical FCD and the FCD of each simulation, to find the value of *G* that minimises the average KS-distance across 40 simulations. This procedure determines the value that allows the model to best simulate the empirical dynamics of functional connectivity in the brain, for each group. (B) Simulated global emergence capacity. (C) Simulated spatio-temporal hierarchy of intrinsic-driven ignition. Every point in this figure is a simulation run. White circle, median; center line, mean; box limits, upper and lower quartiles; whiskers, 1.5x interquartile range. ** $p < 0.01$; *** $p < 0.001$, FDR-corrected.

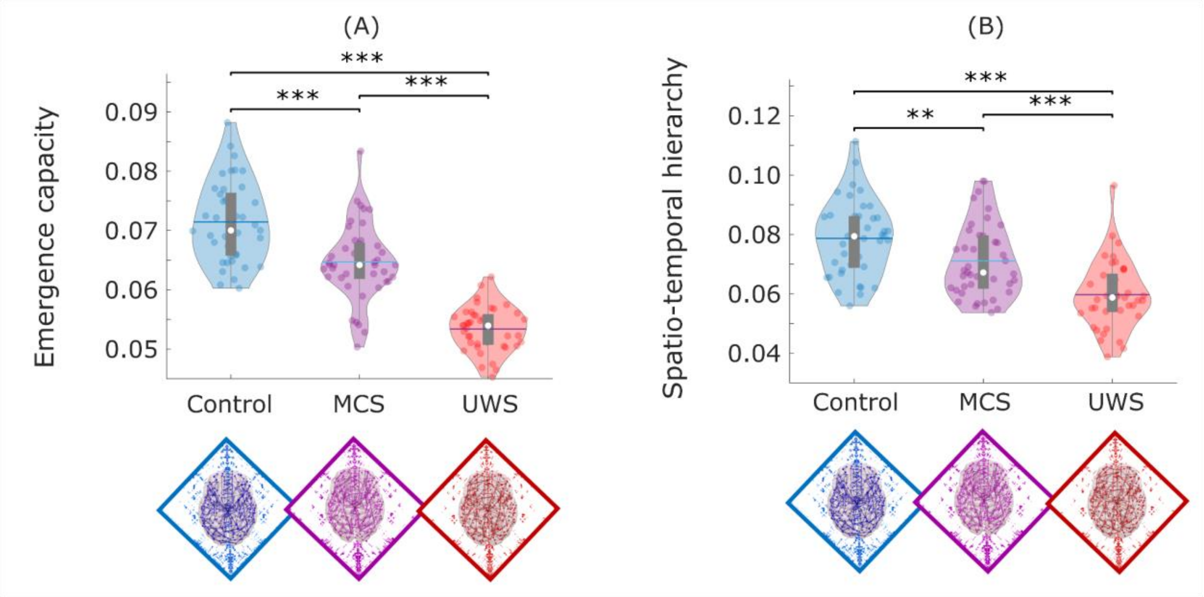


**Figure S7. Replication of whole-brain modelling results with Lausanne-129 parcellation.** (A) Simulated global emergence capacity. (B) Simulated spatio-temporal hierarchy of intrinsic ignition. Every point in this figure is a simulation run. White circle, median; center line, mean; box limits, upper and lower quartiles; whiskers, 1.5x interquartile range. ** $p < 0.01$; *** $p < 0.001$, FDR-corrected.

### Supplementary Tables

**Table S1.** Pairwise statistical comparisons for emergence capacity.

| Contrast | Mean 1 | Mean2 | SD1 | SD2 | tStat | df | EffSize | pVal |
|---|---|---|---|---|---|---|---|---|
| CTRL vs MCS | 0.040 | 0.035 | 0.003 | 0.015 | 3.45 | 28 | 1.25 | 0.002 |
| CTRL vs UWS | 0.040 | 0.031 | 0.003 | 0.003 | 7.24 | 26 | 2.77 | 0.000 |
| MCS vs UWS | 0.035 | 0.031 | 0.05 | 0.003 | 1.90 | 20 | 0.78 | 0.072 |

**Table S2.** Pairwise statistical comparisons for ignition-driven spatiotemporal hierarchy.

| Contrast | Mean 1 | Mean2 | SD1 | SD2 | tStat | df | EffSize | pVal |
|---|---|---|---|---|---|---|---|---|
| CTRL vs MCS | 0.025 | 0.024 | 0.007 | 0.012 | 0.32 | 28 | 0.116 | 0.752 |
| CTRL vs UWS | 0.025 | 0.014 | 0.007 | 0.001 | 4.72 | 26 | 1.806 | 0.000 |
| MCS vs UWS | 0.024 | 0.014 | 0.012 | 0.001 | 2.59 | 20 | 1.066 | 0.018 |

**Table S3.** Pairwise statistical comparisons for whole-brain modal controllability, controlling for DWI acquisition scheme and number of motion-corrupted scans.

| Contrast | Estimate | SE | tStat | EffSize | pVal |
|---|---|---|---|---|---|
| CTRL vs | - | 0.003 | -5.116 | -0.919 | 0.000 |

| | | | | | |
|---|---|---|---|---|---|
| **MCS** | 0.013 | | | | |
| **CTRL vs UWS** | -0.012 | 0.003 | -4.057 | -0.741 | 0.000 |
| **MCS vs UWS** | 0.001 | 0.004 | 0.220 | 0.048 | 0.828 |
